# TnClone: high-throughput clonal analysis using Tn5-mediated library construction and *de novo* assembly

**DOI:** 10.1101/367045

**Authors:** Byungjin Hwang, Sunghoon Heo, Namjin Cho, Duhee Bang

## Abstract

A typical molecular cloning procedure requires Sanger sequencing for validation, which becomes cost-prohibitive and labour-intensive for large-scale clonal analysis of genotype-phenotype studies. Here we present a Tn5-mediated clonal analysis platform TnClone, which uses next-generation sequencing (NGS) to rapidly and cost-effectively analyze a large number of clones. We also developed a user-friendly graphical user interface and have provided general guidelines for conducting validation experiments. Using TnClone, we achieved more than 20-fold cost reduction compared with the cost incurred using conventional Sanger sequencing and detected low-frequency mutant clones (~10%) in mixed samples. We tested our programme and achieved 99.4% sensitivity. Our platform provides rapid turnaround with minimal hands-on time for secondary evaluation as NGS technology continues to evolve.

---

The Sanger method (1) is still regarded as the gold standard for quality control in the analyses of DNA clones (e.g. PCR amplicons or bacterial plasmids). In general, template generated from PCR amplification or direct-cloned plasmid vectors requires a validation step, raising concerns regarding the cost associated with the analysis of a number of plasmid clones. For example, to verify whether the DNA fragment of interest is properly cloned or amplified, Sanger sequencing of the construct in both directions (forward and reverse) is performed. For a 3,000-bp fragment, at least four Sanger sequencing reads are required for accurate validation, which costs at least $12.

For large-insert clones, serial tiling of primers is typically required, which is cost-prohibitive. Mixed ‘dirty’ peaks may confound discerning the true sequence from clone contamination, template heterozygosity [especially insertion and deletions (indels)], incomplete purification, etc. Moreover, Sanger sequencing is not sufficiently sensitive to distinguish mixed peaks when the frequency of the minor contents is <20%. In this case, manual inspection of alignments of a single heterogeneous site can detect the true clone by analyzing high-intensity peaks. However, decomposition of mixed clones with longer inserts and various heterozygous sites poses a challenge.

Advances in next-generation sequencing (NGS) technologies have accelerated the process of analyzing large amounts of DNA sequences while significantly reducing the cost. Recently, there was an effort to analyse plasmid sequence using NGS platform with commercial Nextera reagents (2). However, this method is cost-demanding and adopted robotic system which may not be accessible in general laboratory. In addition, with the explosion of sequencing data, the analysis component is also becoming increasingly important. Moreover, no tool exists to accurately evaluate plasmid DNA clones for biologists who lack computational skills to analyze the data, necessitating an easy-to-use and efficient unified framework.

To overcome these limitations, we present a streamlined platform for analyzing clonality in a multiplexed fashion using an in-house optimized Tn5 transposase-based library preparation (3). The method eliminates expensive and time-consuming steps of making individual sequencing libraries. By taking advantage of combinatorial mosaic end (ME) sequences attached with barcode sequences, we could generate several hundred clonal NGS library in <3 h. In addition, we developed a biologist-friendly python GUI application, TnClone, under Linux/Windows OS for running on a personal computer. Our platform adapts a *de novo* assembly-based approach to assemble long-range insert sequences (i.e. gene of interest) and incorporates the Smith-Waterman (SW) algorithm (4) for reference-based downstream analysis. We expect that our pipeline will aid researchers in analyzing PCR or cloned products and facilitating downstream analysis to provide an NGS-based alternative solution to Sanger sequencing of DNA clones.

## A GUI-based software that makes researcher analysis convenient

TnClone comprises a simple interface, as shown in **Figure 1**. The programme has four major modules: sorting, trimming, assembly, and downstream analysis. Sorting is processed by searching for ME and Tn5 barcode sequences from raw NGS data. This step automatically demultiplexes the raw fastq files using barcode combinations included in the library preparation step. At the trimming step, we used Trimmomatic-v0.36 (5), with default options to remove low-quality data. Subsequently, a de Bruijn graph-based assembly takes input from previously trimmed NGS data. Users can optionally skip the trimming step and directly jump to perform a *de novo* assembly. Then, alignment is performed using the SW algorithm. Because the exhaustive SW search is too slow, we used a Stripped SW algorithm(6) to accelerate alignment steps if insert sizes were large. Finally, TnClone will give users two types of reports: i) a simplified variant call format and ii) a brief summary file containing information of DNA/Protein error/error free and the number of contigs assembled.

**Figure 1.**
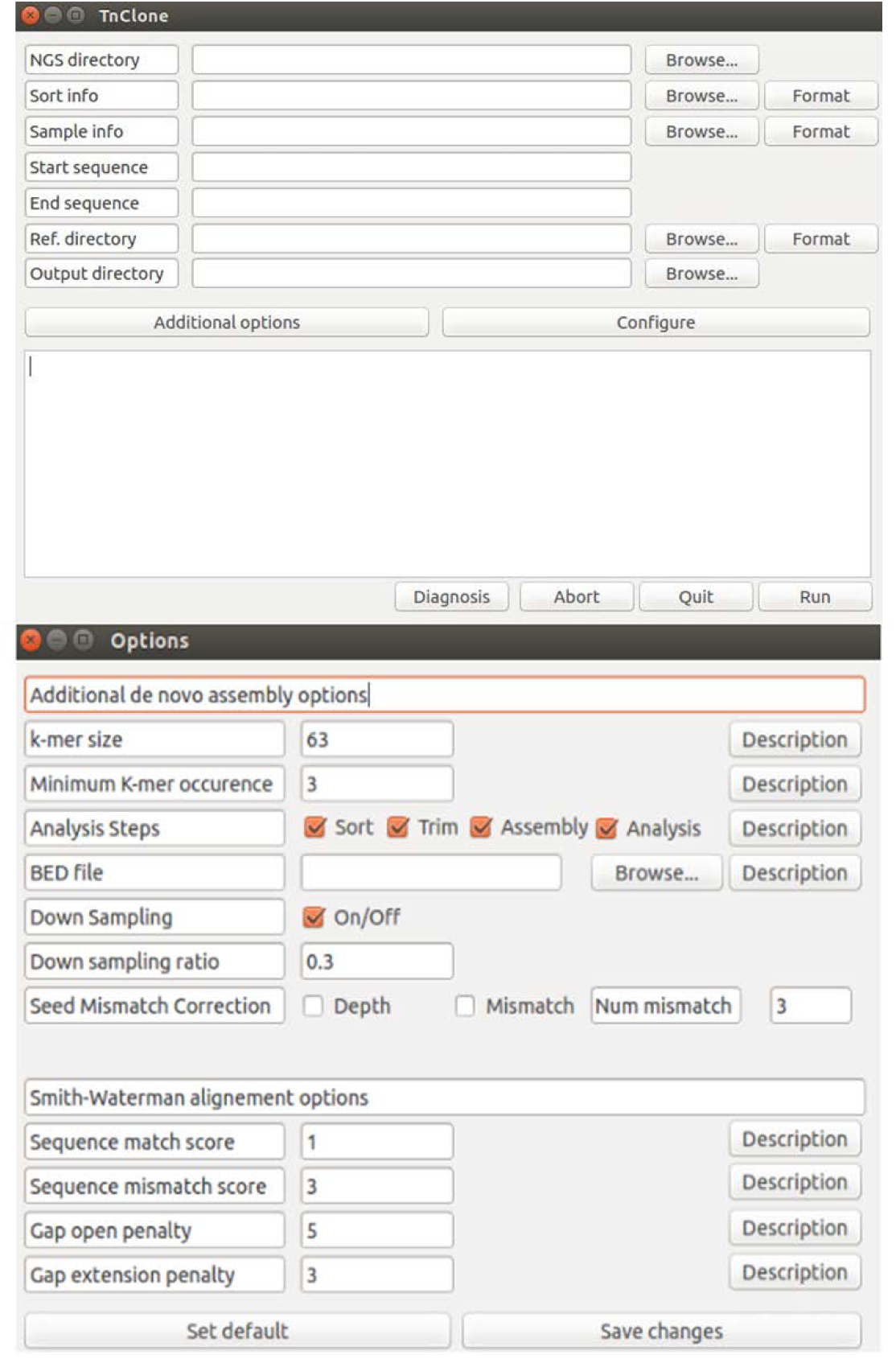
TnClone’s interface. TnClone comprises two windows: the main analysis window (upper) and the option selection window (lower). The main window shows the general inputs for TnClone. All fields are required except for the Start sequence and End sequence fields. In the options window, users can change the value of parameters required for downstream analysis. The default parameters of the TnClone software are shown below. These parameters are in-house tested for optimal results.

Users can control several features. In the main window (**Figure 1**), users can choose the start (seed) and end sequences for assembly. All fields except these two fields are required. On the additional options screen (**Figure 1**), users can tweak assembly parameters and alignment options for downstream analysis. On this screen, if users do not specify seeds and end sequences for assembly, users must specify a region file (BED file field) to automatically detect seeds and end sequences for assembly. Users can change *k*-mer (consecutive k-length nucleotides) size to control the accuracy of the assembly and the minimum *k*-mer occurrence to pre-remove *k*-mers not exceeding this value (often considered to be erroneous *k*-mers). If the file size is too large, then one can downsample the original data and perform assembly. Also users can set a maximum of three mismatches for initial seeds. If one has a reference sequence for downstream analysis, TnClone provides overall coverage information (by checking if certain regions are not sequenced) on clicking the ‘Diagnosis’ button (see **Figure 1** and **Supplementary Figure 2**). For samples with a reference sequence, Alignment Options are required for downstream analysis. We set our default parameters from in-house tested optimized values.

**Figure 2.**
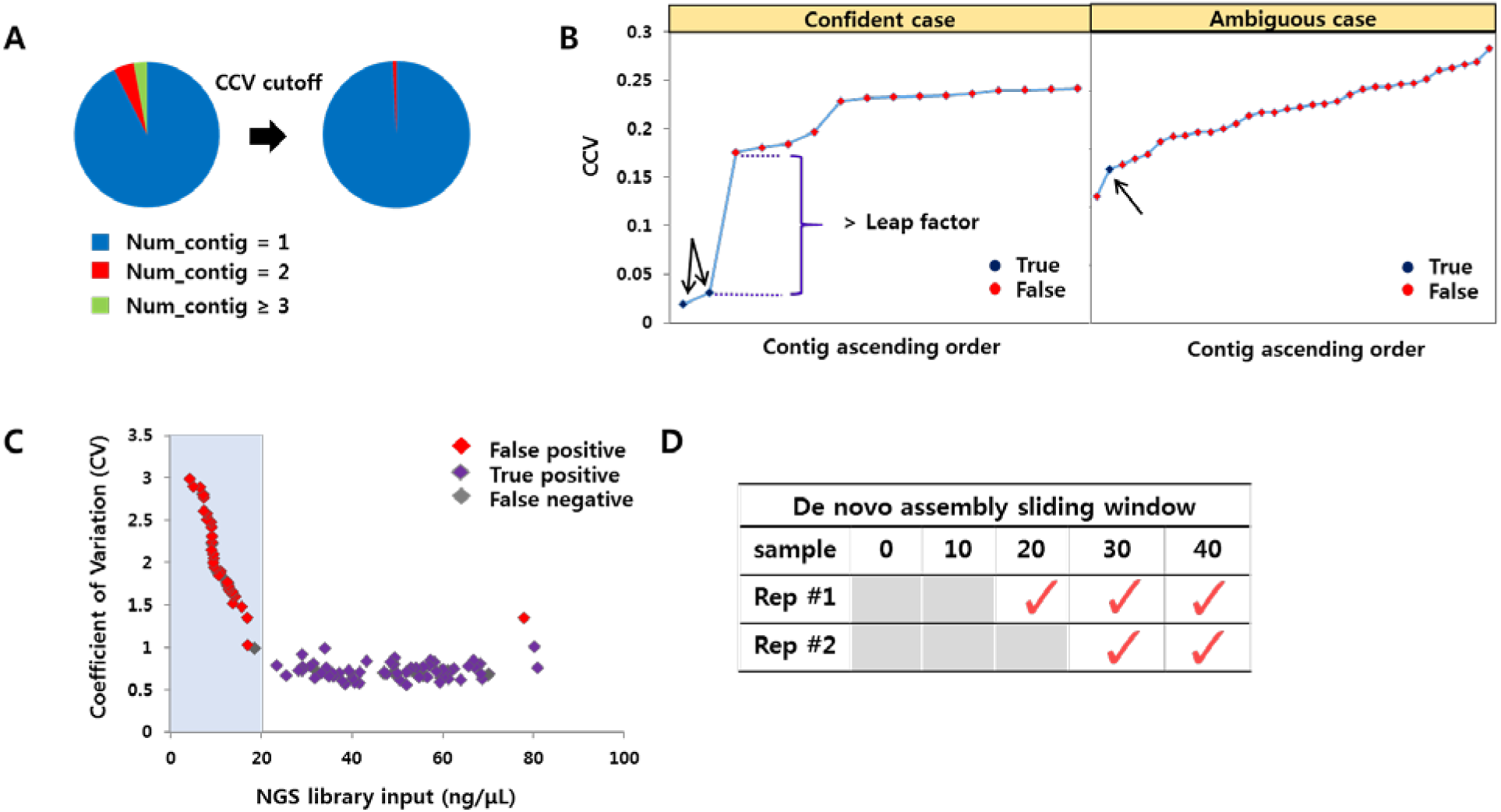
Analysis of clonal information using the TnClone software. (**a**) Pie chart for assembled contigs of initially assembled candidate contigs and following the final selection step. (**b**) Knee plot of the coefficient of variation scores of candidate contigs for representative samples. The black arrow indicates the true contigs validated by Sanger sequencing of the pure cultured colonies. If the leap factor between two consecutive points (i, I + 1) exceeds a certain value, ≤i^th^ contigs are considered as true contigs. On the right, the ambiguous case is shown with no clear leap point. (**c**) The correlation between the next-generation sequencing (NGS) library input versus coefficient of variation of scFv clones is shown (n = 91). The lower the input DNA available for NGS sequencing, the more variance exists in the depth distribution of the assembled contigs, leading to more false contigs. (**d**) Analysis of PCR amplicons using TnClone. We used a sliding window approach to detect a reliable assembly for two replicate experiments and found that a 30-bp flanking position is required for accurate assembly.

## *De novo* assembly of cloned constructs of *cas9* and scFv

To assess whether TnClone can resolve specific variants in a single clone, we tested 801 samples, 144 of which were randomly verified with Sanger sequencing (scFv and Cas9 clones), and a sensitivity of 99.4% was achieved. With respect to the 801 analyzed clones, the majority (93%) of the initially assembled contigs were single contigs (**Figure 2a**). When we further reduced the true candidates using a contig coefficient of variation (CCV, see the **Supplementary Methods** section) score cut-off, unique contigs were highly enriched (99%), implying that the clone contamination rate was low. Notably, we reliably detected dual clones by applying a CCV score from multiple, initially assembled candidate contigs (**Figure 2b left**). However, we found one case with no clear leap point, as shown in a 1:1 mixture experiment (**Figure 2b right** and **Supplementary Figure 4**). For the case with 1:1 mixture, amplification or sequencing error might have caused bias in the depth distribution at the polymorphic site. We defined this case as ‘ambiguous’ in contrast to the ‘confidence’ case, in which we can clearly separate the true contigs using cut-off scores. To offer a practical guideline for the minimum input plasmid DNA for contig detection, we evaluated scFv clones and classified false and true positive ones. At a 20-ng/μl cut-off, we could detect most of the true positive clones (~99%), which showed a lower coefficient of variation than the false positive ones (**Figure 2c**).

## Mixed sample analysis

To demonstrate whether TnClone could distinguish mixed clones, we first performed three replicates of a mixing experiment containing the mixture ratios of 1:1, 2:1, 3:1, and 9:1. For comparison, we performed simulation analysis with a synthetic library. Synthetic libraries were generated using the ART simulator, as described in the **Supplementary Methods** section. Using the four dilution conditions, we could reliably detect variant contigs with a frequency as low as 10%, and we observed a strong correlation between real and simulated data (**Supplementary Table 1** and **Supplementary Figure 5**).

## Assembly of the cloned product directly from bacterial lysates

To evaluate whether Tn5 integration directly into cells from bacterial colonies can reduce intervening clean-up steps, we tested five genomic libraries (cloned plasmid DNA + genomic DNA mixture, **Supplementary Methods**) and set TnClone default parameters for assembly. After mapping assembled contigs to the reference sequence, TnClone achieved alignment identity of >99% on average for five plasmid DNA sequence, suggesting high-quality *de novo* assembled contigs.

## Analysis of PCR amplicons

Next, we used TnClone to test whether products from PCR experiments could be assembled. Samples were amplified directly from cloned vectors (pCMV-BE3, AddGene accession #73021) 759 bp, 1398 bp, and 2126 bp in length. NGS libraries were sequenced using the Tn5 tagmentation method, similar to that in the plasmid library preparation described above. We found that the sequencing depth was distributed in a Gaussian pattern, which prohibits full-length assembly of the amplified product. This was likely due to insufficient sequencing depth at both ends (5' and 3' ends) of PCR amplicons. Using a sliding window approach (**Figure 2d**), we concluded that the lowest required length for the assembly was 30 bp farther from the original reference sequence. This indicated that some flanking sequence was required for both ends to reliably assemble regions of interest.

## DISCUSSION

We developed TnClone, a high-throughput platform for analyzing DNA clones. TnClone focuses on utilizing a GUI to make a user-friendly platform for general biologists. To our knowledge, this is the first programme for a high-throughput NGS system aimed at large-scale clone validation. By simple clicking and scrolling, users can analyze sequences from scratch without processing the intermediate files from raw data individually. In addition, we developed a custom Tn5 transposase with index sequences attached to ME sequences. This allows further multiplexing of the samples after adding Illumina’s index bases for sample identification. In addition, library preparation is simple, does not require specialized experimental techniques, and takes <3 h. Notably, compared with conventional Sanger sequencing, the entire cost for analyzing a clone is 20-fold (**Supplementary Table 2**) less expensive.

In summary, our streamlined Tn5 transposase-based NGS library preparation and *de novo* assembly-based clonal analysis facilitates verification of large numbers of clones in a high-throughput manner. As the scalability of biological experiments increases, we expect that an NGS-based clonal analysis platform will be broadly applicable to various validation experiments.

## Accessibility

TnClone is available at https://github.com/tahuh/tnclone. The source code is written in python, and the source package is distributed under a GPL licence (v3).

## ACKNOWLEDGEMENTS

We would like to thank the Bang’s laboratory members for their helpful discussions on the manuscript and for testing the algorithm. We would also like to thank Junho Jung’s laboratory members for kindly donating cloned plasmid vectors of single-chain antibody. This work was supported by Pioneer Research Center Program (NRF-2012-0009557), Mid-career Researcher Program (2015R1A2A1A10055972), and Bio & Medical Technology Development Program (NRF-2016M3A9B6948494) through National Research Foundation of Korea funded by the Ministry of Science, ICT & Future Planning. This research was partially supported by the Graduate School of YONSEI University Research Scholarship Grants in 2017.

## AUTHOR CONTRIBUTIONS

B.H., S.H. and N.C performed the experiments. B.H. and S.H. analysed the data and wrote the paper. D.B. supervised the project.

---

Graphical Table of Contents

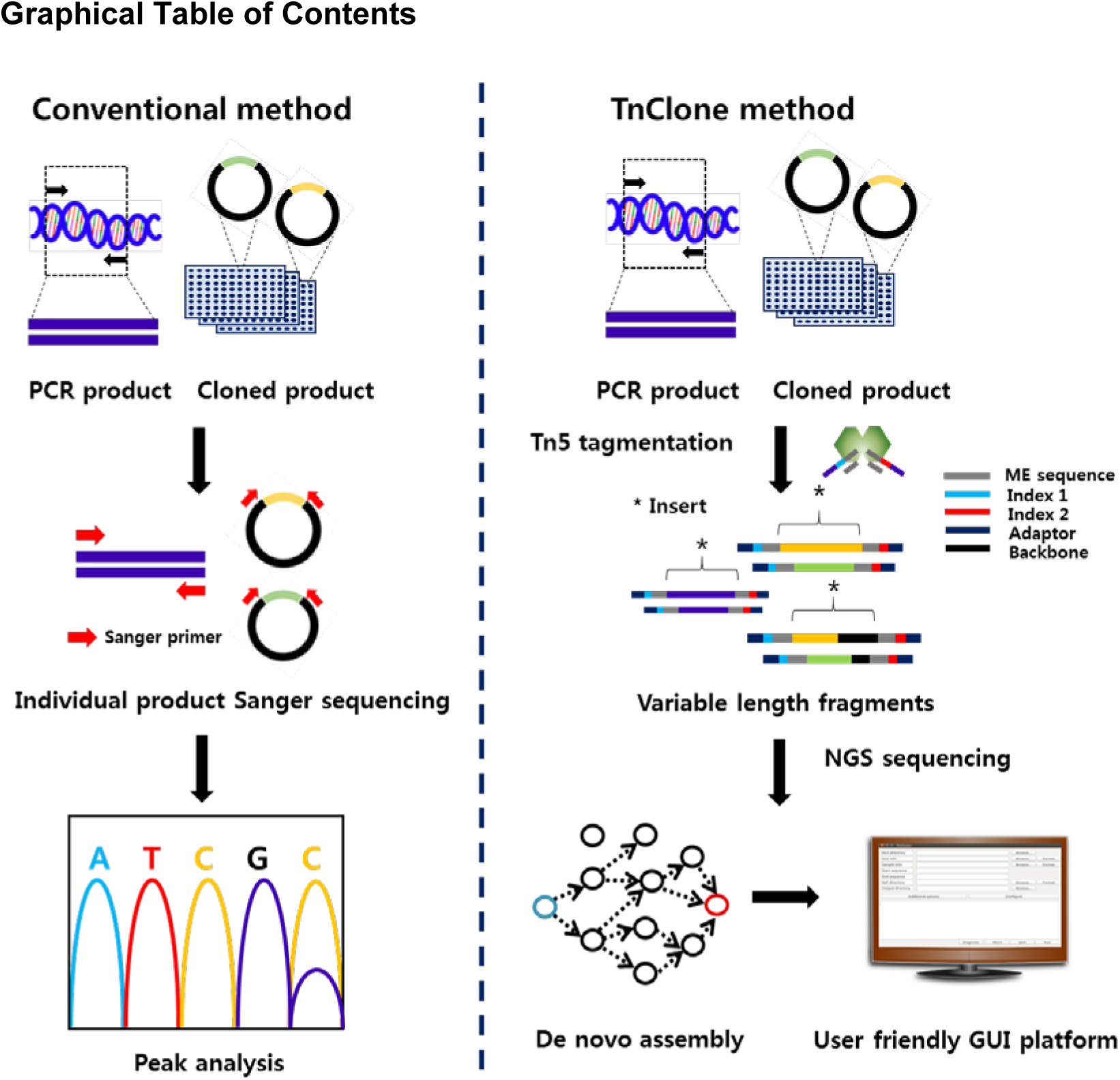

